# Transdermal, but not oral mucosa exposure, causes sublethal leptospirosis in mice

**DOI:** 10.1101/334219

**Authors:** Nisha Nair, Maria Gomes-Solecki

## Abstract

The goal of this study was to determine the timing of *Leptospira* dissemination after infection using natural physiologic routes and its correlation with signs and symptoms of leptospirosis. Groups of C3H-HeJ mice were sub-lethally infected through exposure of a dermis wound and the oral mucosa to *L. interrogans* serovar Copenhageni, and compared to mice infected via standard laboratory practice of intraperitoneal inoculation. We found that transdermal infection leads to delayed bacterial dissemination in blood and urine when compared to intraperitoneal infection using the same infectious dose, and that the overlapping presence of *L. interrogans* in both fluids was twice as long as the standard intraperitoneal infection. Furthermore, dissemination of *L. interrogans* and disease did not occur in mice infected with the same dose of *L. interrogans* via the oral mucosa. Over the course of these studies we also observed that the timing of exit of *Leptospira* from blood and establishment of colonization of the kidney in the second week of infection correlated well with weight loss, but not with hypothermia. Thus, a non-invasive sign of leptospirosis such as weigh loss can be used to monitor bacterial dissemination and disease stage in mice. Our findings underline the importance of precise determination of windows of pathogen dissemination in biological fluids and how the route of infection affects the progression of disease. These studies could help focus the development of better treatment strategies and new technologies for diagnostic use.

## INTRODUCTION

Zoonotic diseases are a major concern to human health even in the current modern era of medical and scientific advancement. Leptospirosis, caused by pathogenic *Leptospira sp*. is a neglected emerging zoonotic disease prevalent in industrialized urban, suburban, and rural regions, and is endemic to areas with tropical and temperate climate. It is well established that pathogenic *Leptospira sp.* have a wide range of vertebrate animals as its reservoir host, most of which are asymptomatic carriers. Humans are incidental hosts. Rodents, specifically rats and mice, are the main carrier hosts that contaminate water and soil with their urine (1). Humans acquire infection after exposure to contaminated sources through cuts and scrapes in the skin and mucous membranes or consumption of contaminated food. Symptoms can range from asymptomatic to mild febrile illness culminating in multi-organ failure, if left untreated (2).

Our current understanding of *Leptospira sp*. pathogenesis and virulence factors derives from studying laboratory animals, specifically the acute state of infection through intraperitoneal inoculation of hamsters (3), (4), (5), (6), (7), (8). Rats have been used to study chronic infection (9), (10). Mice have been used to evaluate acute lethal and sublethal infections (11), (12), (13), (14), (15), (16). Exposure to Leptospira under physiological conditions, i.e. the penetration of *Leptospira* through skin and mucosa, was recently evaluated in rats (17) and in mice (18).

In this study we used the sublethal mouse model to evaluate dissemination of *L. interrogans* after inoculation using two natural physiological routes (oral mucosa and transdermal) compared to the standard laboratory practice of intraperitoneal infection.

Our goal was to correlate clinical signs disease progression after natural infection with pathogen burden in biological fluids and with measurable biomarkers of disease severity.

## MATERIALS AND METHODS

### Bacterial strains

We used *Leptospira interrogans* serovar Copenhageni (LIC) strain Fiocruz L1-130, originally isolated from a patient in Brazil and passaged in hamster (passages number 3 and 4). *L. interrogans* was cultured as previously described (18) and enumerated by dark-field microscopy (Zeiss USA, Hawthorne, NY), and by StepOne Plus qPCR system (Life Technologies, Grand Island, NY).

### Animals

Female, 10-week old, C3H/HeJ mice were used for transdermal and intraperitoneal studies. Mice were purchased from The Jackson Laboratories (Bar Harbor, ME) and acclimatized for one week at the pathogen-free environment in the Laboratory Animal Care Unit of the University of Tennessee Health Science Center.

### Ethics statement

This study was carried out in accordance with the Guide for the Care and Use of Laboratory Animals of the NIH. The protocols were approved by the University of Tennessee Health Science Center (UTHSC) Institutional Animal Care and Use Committee, Animal Care Protocol Application, Permit Number 14-018.

### Infection of mice

Intraperitoneal infection was done as described previously using a dose of ∽10^8^ *L. interrogans* serovar Copenhageni FioCruz in sterile PBS. For transdermal infection, a wound was generated on the back of anesthetized mice. One square inch area on the lower back was shaved and the exposed skin was scraped with a sterile razor just enough to create a superficial wound (no bleeding). Subsequently, 50-100 μl of *L. interrogans* culture (∽10^8^ cells) was applied on the transdermal wound and covered using an occlusive bandage. The protective bandage was removed the next day. For oral mucosa infection, the same amount of *L. interrogans* were deposited in the buccal cavity. Groups of mice inoculated with endotoxin free PBS (Dulbecco) into the peritoneum (IP Ctrl), into the buccal cavity (OM Ctrl) and deposited on the transdermal wound (TD Ctrl) were kept as negative controls. Body weight and temperature (°C) were monitored daily and urine was also collected on a daily basis for 15 days post infection. Blood (up to 20 μl) was collected every other day by tail nick for 15 days post infection. At termination, kidneys were collected for quantification and culture of spirochetes, and for quantification of inflammatory transcripts. Spleen from mice infected via transdermal route was also collected at termination and processed for FACS analysis as described (15).

### ELISA

Concentration of immunoglobulin IgM, IgG, IgG1, and IgG2a in mouse serum was determined using Ready-Set-Go ELISA kits (eBioscience). Leptospira-specific-IgM and - IgG antibodies were detected in serum of infected mice against heat-killed *Leptospira* coated on microplate wells at a concentration of 4mg/ml.

### RT-PCR, and q-PCR

DNA was extracted per the manufacturers’ instructions from urine, blood, and kidney using a NucleoSpin tissue kit (Clontech). Quantification of *Leptospira* was done using TAMRA probe and primers from Eurofins (Huntsville, AL) by real-time PCR (qPCR) (StepOne Plus). RNeasy mini kit (Qiagen) was used to extract total RNA followed by reverse transcription using a high-capacity cDNA reverse transcription kit (Applied Biosystems). Real-time PCR on the cDNA was performed as described (15). For RT-PCR, we used TAMRA probes specific for inducible nitric oxide synthase (iNOS), Collagen A1 (ColA1), keratinocyte-derived chemokine (KC, CxCL1), macrophage inflammatory protein 2 (MIP-2, CxCL2), RANTES (CCL5), tumor necrosis factor alpha (TNF-α) and interferon gamma (IFN-γ). β-actin was used as control for the comparative CT method (19).

### Kidney histopathology

Kidneys were excised from mice after euthanasia and fixed in 10% formalin. Presence of Leptospira in the kidney was determined by Whartin-Starry silver staining of kidney sections at termination. Formalin-fixed paraffin embedded tissues were stained with periodic acid-Schiff – Diastase (PAS-D). Stained sectioned were evaluated for interstitial inflammation, glomerular morphology and size using an Axio Zeiss Imager A1 light microscope.

### Processing of spleen

Spleens were excised aseptically after euthanizing the mice and transported in RPMI-1640. Each sample was processed individually to prepare single cell suspension by teasing the tissues gently in between two sterile frosted slides in Hanks balanced salt solution (Cellgro, VA). ACK lysing buffer (Lonza, MD) was used to lyse red blood cells and washed with RPMI-1640 with 10% fetal bovine serum and 0.09% sodium azide. Single cells were prepared by passing the cell suspension through 70 μm- and 40 μm-pore-size nylon filters (BD Falcon, MA) sequentially. Cells were counted and tested for viability, using acridine orange/propidium iodide staining, on Luna-FL automated cell counter (Logos Biosystems, South Korea).

### Immunophenotyping

Single cell suspension from spleen were stained by the following protocol: cells were incubated in Fc block solution (Tonbo Biosciences) for 15 min at 4°C prepared in staining buffer (Ca2+ and Mg2+ free PBS containing 3% heat-inactivated fetal bovine serum, 0.09% sodium azide, 5 mM EDTA). The cells were washed twice and incubated with appropriate surface markers in the dark for 30 min at 4°C. The lineage surface markers used for the study are described in (15) and (19). LSR II flow cytometer (BD Immunocytometry Systems, CA) equipped with 405nm, 488nm, 561nm, and 640nm excitation lasers was used to acquire surface stained cells on BD FACS Diva software (BD Biosciences). Data was analyzed on FlowJo software (TreeStar, OR). In order to distinguish positive cell population from negatively stained cells, gating using Fluorescence-minus-one (FMO) controls was used. Single color beads were prepared from BD Comp Beads (eBiosciences) for compensation. 100,000 events were acquired per sample.

### Statistics

Two-tailed unpaired t-test with Welch’s correction was used to analyze differences between infected and non-infected groups in body weight, body temperature and T cell populations. Ordinary one-way ANOVA was used to evaluate differences in inflammatory and fibrosis mediators between all groups: Ctrl, IP, OM, TD. Statistical analysis was done using GraphPad Prism software, α = 0.05.

## RESULTS

### Mice infected through the oral mucosa do not lose weight, unlike mice infected through the transdermal route

10-week-old female C3H/HeJ mice were infected with 10^8^ *Leptospira interrogans* Copenhagani FioCruz by deposition of inoculum in the buccal cavity or on a transdermal wound (TD). Groups of mice inoculated with the same dose of *L. interrogans* into the peritoneum (IP) were kept as positive controls. Groups of uninfected mice were kept as negative controls. Weight records kept for 15 days (Fig. 1A) post infection showed that mice infected via OM did not lose weight, and temperature records (Fig. 1B) of infected mice were not different from controls. However, in mice infected via the TD route differences between the weight records were significant between infected and non-infected groups, although temperature did not change between the two groups. As expected, mice infected via the IP route lost significant amounts of weight and temperature in the two weeks following infection.

**Figure 1.**
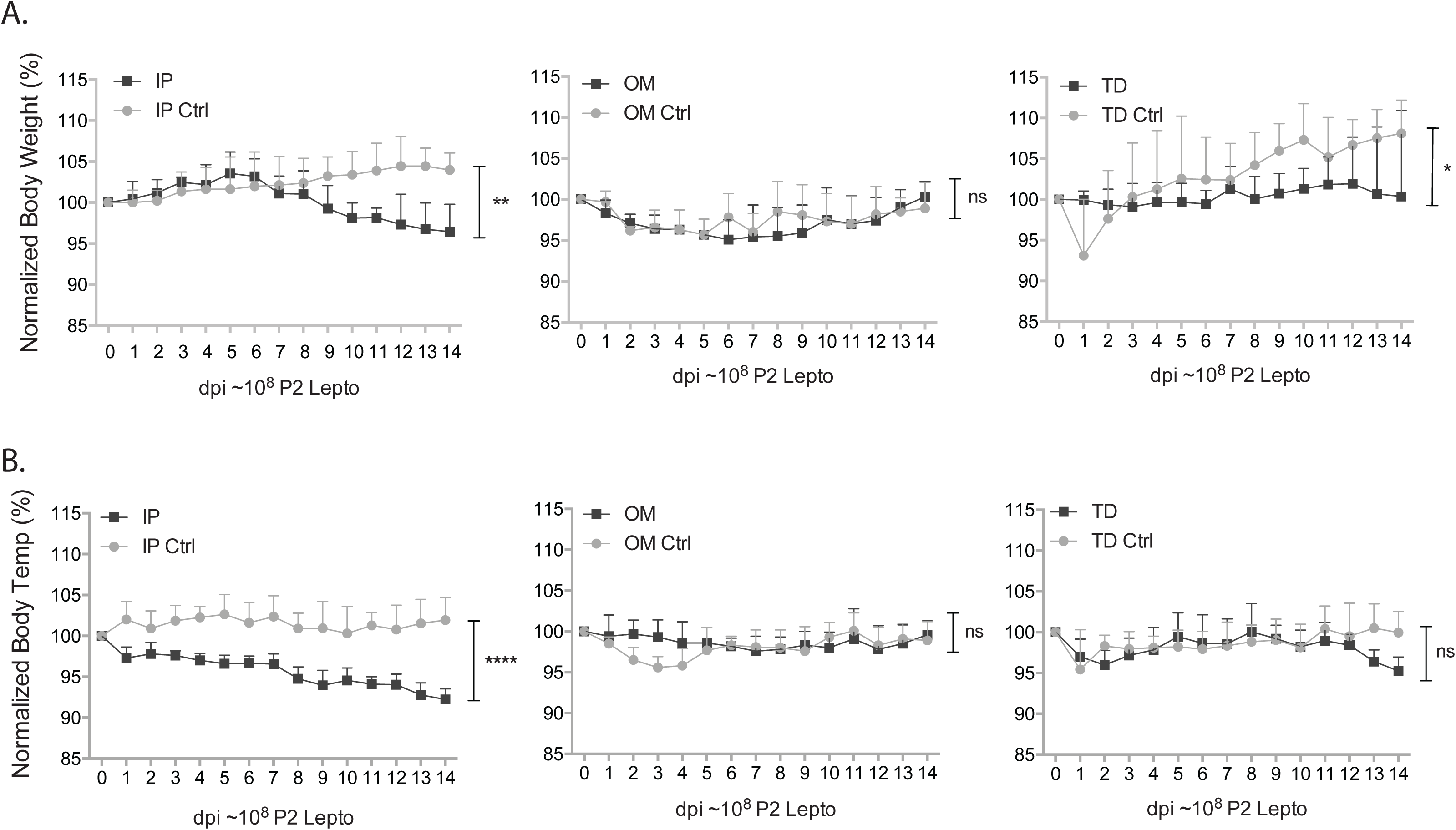
Signs of disease progression after infection via two natural physiologic routes: weightloss and temperature. Groups of C3H-HeJ mice were inoculated with an equivalent high sublethal dose of *L. interrogans* serovar Copenhageni FioCruz (10^8^) via oral mucosa (OM) and transdermal (TD) routes of infection and compared to standard laboratoy practice intraperitoneal inoculation (IP). Body weight (A) and rectal temperatures (B) were recorded daily for two weeks post infection. Statistics by Unpaired t-test with Welch’s correction, weightloss IP p=0.0049, OM p=0.3459 and TD p=0.0296; temperature IP p<0.0001, OM p=0.1619 and TD p=0.2161. N=4 mice per group. Data represents one of three (IP and TD) and two (OM) independent experiments.

### Mice infected through the transdermal route have high levels of *L. interrogans* in blood and urine, unlike mice infected through the oral mucosa

The kinetics of *L. interrogans* dissemination in blood (Fig. 2A) was considerably different between the three routes of infection. Mice infected with 10^8^ spirochetes via the TD route had a maximumof 10^6^ *L. interrogans* per ul of blood for about 1 week between days 6-13 post infection, whereas mice infected via direct inoculation of 10^8^ spirochetes into the peritoneum cavity had about 10^2^-10^3^ spirochetes per ul of blood between days 1-7 post infection. *L. interrogans* did not disseminate in blood and urine of mice infected with the same dose via oral mucosa. Regarding colonization of the kidney (Fig. 2B), *Leptospira* was detected in urine 5 days after it was first detected in blood after IP infection (on day 6), and 3 days after it was first detected in blood after TD infection (on day 9). The period when we detected *L. interrogans* both in blood and in urine was twice as long for TD (5 days of overlap) than for IP infection (3 days of overlap). After establishment of kidney colonization, mice shed about the same amount of Leptospira in urine (∽10^5^/μl) following IP and TD infection (Fig. 2B).

**Figure 2.**
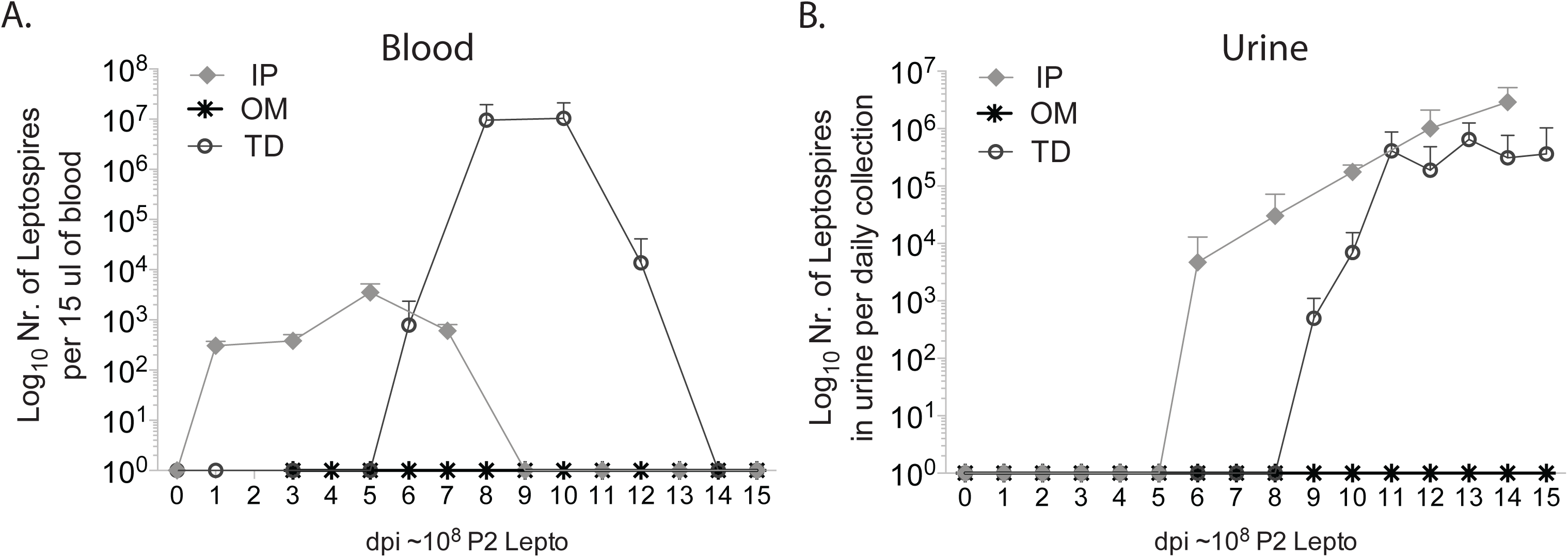
Leptospira dissemination in blood and urine. Groups of C3H-HeJ mice were inoculated with an equivalent sublethal high dose of *L. interrogans* serovar Copenhageni FioCruz (10e8) via two natural physiological routes of infection (oral mucosa (OM) and transdermal (TD)) as compared to standard intraperitoneal inoculation (IP). Bacterial burden in blood (A) and urine (B), was determined by 16s rRNA qPCR. N=4 mice per group. Data represents one of two (IP v OM) and three (IP v TD) independent experiments.

### Kidneys from infected mice had equivalent amounts of Leptospira independently of route of infection (IP or TD) and showed histopathology signs of inflammation

Presence of Leptospira in the kidney was determined by Whartin-Starry silver staining of kidney sections at termination. In the infected groups (TD and IP), spirochetes appeared as black colored aggregates and as dispersed single cells interspersed through the renal tissue, which was absent in the uninfected control (data not shown). Histopathology analysis of PAS-D stained paraffin embedded kidney infected tissue (TD and IP) showed increased cellular infiltration and the glomerular size was reduced by one-third as compared to controls as observed in (15). We detected the same amount of *Leptospira* per mg of kidney tissue (5×10^5^) in IP and TD infected mice, whereas no spirochetes were detected in controls or in mice infected via the oral mucosa (OM) (Fig. 3A). We confirmed the viability of the spirochetes quantified in the infected kidney, by culturing the tissue at 30° C in EMJH. Samples collected from cultures at d0, d3 and d8 post kidney tissue inoculation (d0) showed a growth curve of increasing numbers of cells for tissue collected from mice infected via IP and TD but not for tissue harvested from controls or mice infected via OM (Fig. 3B).

**Figure 3.**
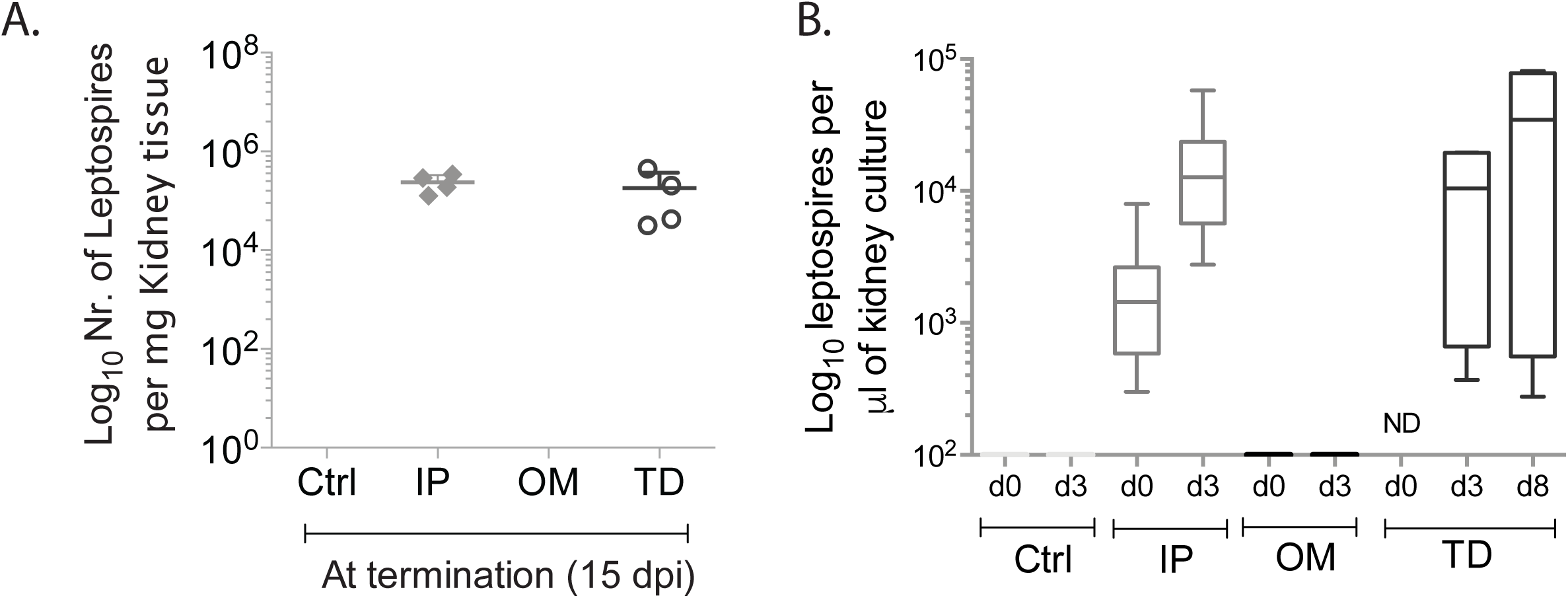
Leptospira burden and viability in kidney tissue. Groups of C3H-HeJ mice were inoculated with equivalent sublethal high doses of *L. interrogans* (10^8^) via two natural physiological routes of infection, oral mucosa (OM)) and transdermal (TD) compared to standard intraperitoneal (IP). Leptospira burden and viability in kidney tissues was determined by qPCR of 16s rDNA from kidney tissue and from kidney culture. ND, Not determined. N=4 mice per group. Data represents one of two (IP v OM) and three (IP v TD) independent experiments.

### Transcription of pro-inflammatory mediators and fibrosis markers in kidney following transdermal (TD) and oral mucosa (OM) infection

Analysis of pro-inflammatory mRNA showed significant increases of innate response mediators (CxCL1/KC, CxCL2/MIP-2, CCL5/RANTES, TNF-α) and Th1 IFN-γ in kidneys from mice infected via TD and IP but not in mice infected via OM or from controls (Ctrl). Differences for all pro-inflammatory markers were statistically significant (Fig. 4). Analysis of inflammatory and fibrosis mediators, inducible nitric oxide (iNOS) and fibroblast activation marker collagen A1 (ColA1), were not different between kidneys from mice infected via IP, OM and TD or from controls (Ctrl).

**Figure 4.**
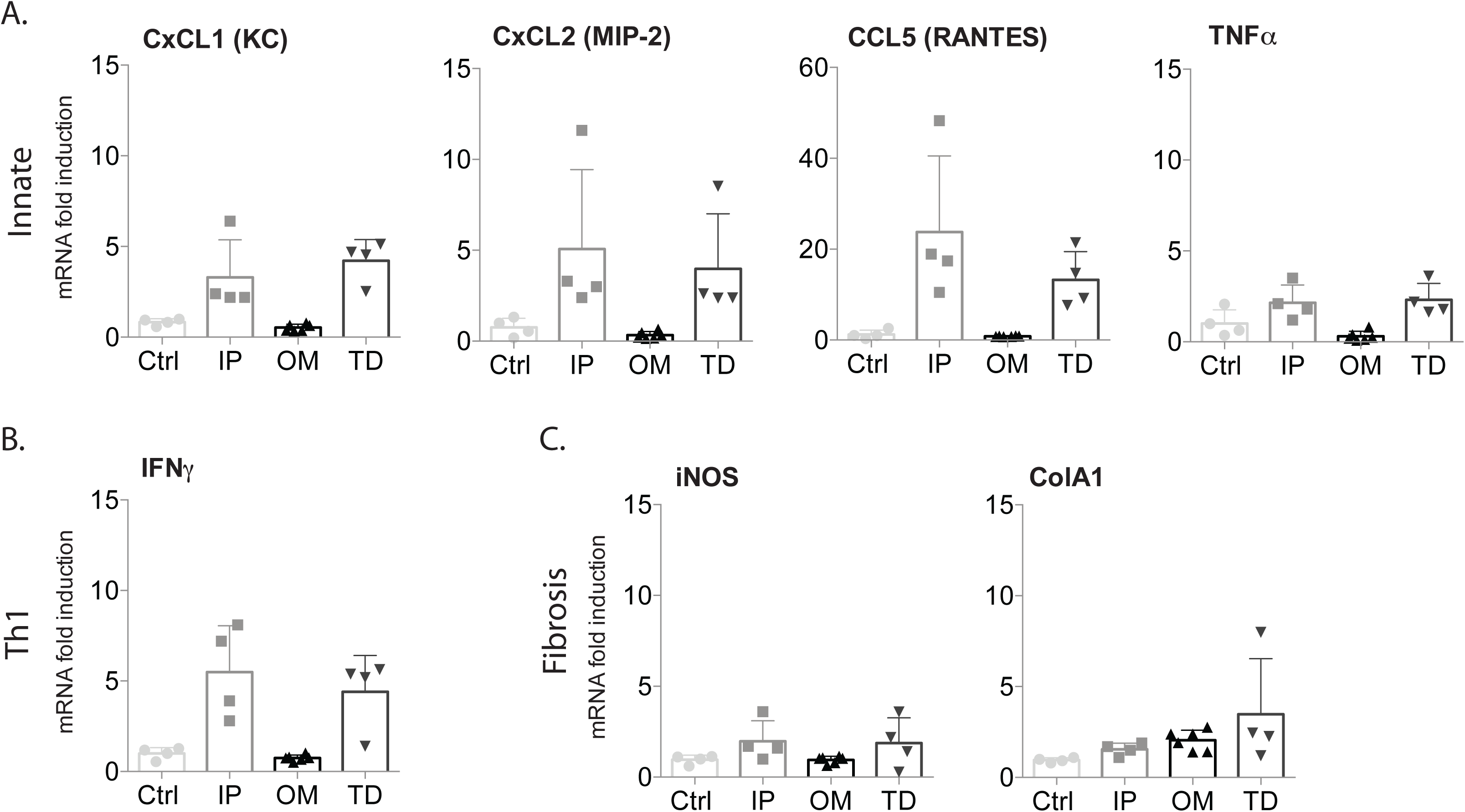
Transcription of pro-inflammatory and fibrosis markers in kidney of mice after inoculation with *L. interrogans* using physiological and standard routes of infection.C3H-HeJ mice were infected with equivalent sublethal high doses of *L. interrogans* serovar Copenhageni (10^8^) and kidney was collected two weeks post infection for quantitative PCR analysis of reverse-transcribed mRNA. A. Innate response chemokines CxCL1, CxCL2 and CCL5 and cytokine TNF-alfa; B. Th1 cytokine IFN-gamma and C. inflammatory and fibrosis markers inducible nitric oxide (iNOS) and fibroblast activation marker collagen A1 (ColA1). Statistics: Ordinary One-Way ANOVA between IP, OM and TD: KC p=0.0017; MIP-2 p=0.0478; RANTES p=0.0090; TNFa p=0.0016; IFNg p= 0.0024; iNOS p=0.2069; ColA1 p= 0.2654. N=4 per group. Datarepresents one of two (IP v OM) and three (IP v TD) independent experiments.

### B and T cell response to *L. interrogans* following transdermal infection

Quantification of total immunoglobulin (Ig) by ELISA showed ∽1358 μg/ml of IgM and ∽ 1221 μg/ml IgG which was isotyped into 722 μg/ml of IgG1 and 144 μg/ml of IgG2a. We confirmed that increased concentrations of immunoglobulins were *Leptospira* specific by testing the same serum against heat-killed *Leptospira* coated on ELISA plates (Fig. S1A). Analysis of T cell populations isolated from spleen from infected (TD) and uninfected (Ctrl) mice showed that although there was no significant increase of CD4+ splenocytes incomparison to the wound control there was a reduction of CD8+ cytotoxic T cells (Fig. S1B). A further breakdown of the T cell populations in spleen was done using CD44 and CD62L surface markers. An increasing expression of CD44 and a declining expression of CD62L on the cell surface of CD4+ and CD8+ indicated activation of the T cells from naϊ ve to effector. There was no difference in numbers of memory T cells, which is represented by increased expression of CD62L surface markers. These results are consistent with the T cell immunophenotypes we previously determined after infection of mice via IP route [8].

## DISCUSSION

One of our goals is the development of mouse models of Leptospirosis that accurately recapitulate disease progression following infection via natural transmission routes. Knowing accurate timing of bacterial dissemination after natural infection and correlation with disease symptoms helps focus the development of better treatments and new strategies for diagnostic use.

We characterized a mouse model of sublethal infection using C3H-HeJ inoculated with 10^6^ *L. interrogans* serovar Copenhageni FioCruz after 10 weeks of age, via the standard laboratory practice intraperitoneal route (15). As follow up, we tested a natural enzootic route of infection (conjunctival, CJ) and we found that a much higher dose of spirochetes (∽10^8^) was necessary to produce disseminated infection (18). In the study reported here, we evaluated how natural exposure of C3H-HeJ to *L. interrogans* ser.

Copenhageni via two physiologic routes recapitulate Leptospirosis progression as compared to the standard laboratory practice of intraperitoneal infection. We used equivalent doses of the same pathogenic serovar of *Leptospira* for transdermal, oral mucosa and intraperitoneal inoculations. We found that transdermal inoculation of *L. interrogans* resulted in bacterial dissemination in blood which was followed by kidney colonization and shedding in urine as observed previously for IP (15) and CJ (18) infections, whereas infection through the oral mucosa did not result in bacterial dissemination or tissue colonization. Of note, the side by side comparison of IP and TD infections using the same infectious dose (∽10^8^) resulted in distinct timings of bacterial dissemination, with IP infection leading to immediate dissemination in blood (d1-d8) and TD infection leading to dissemination 6 days post infection (d6-d13), which was similar to the timing observed after infection with 10^8^ *Leptospira* through the conjunctival route (18). Colonization of the kidney and shedding in urine happened in the second week of infection in all cases. Furthermore, the overlapping window between the two phases (blood dissemination and urine shedding) in mice infected transdermally was nearly twice as large as the observed for IP infection using the same dose, which is consistent with our observation when we used another physiologic route of bacterial infection, the conjunctival route (18). Over the course of three studies ((15), (18) and current) we also observed that the timing of exit of *Leptospira* from blood and establishment of colonization of the kidney in the second week of infection correlated well with weight loss, but not with hypothermia, independently of route of infection. Thus, we are confident that weight loss can be used to monitor pathogenic *Leptospira* dissemination and disease stage in mice.

When we analyzed the kidneys of infected (TD) mice with PAS-D stain we observed reduced glomerular sizes and higher immune cell infiltration as well as moderate interstitial nephritis in the infected kidneys than in the wound control. Silver staining showed the presence of healthy coiled *L. interrogans* clusters within the renal tubules, which were proven viable after culture from the kidneys into EMJH medium. These results were not different than our observations using the IP (15) and the conjunctival (CJ) infection (18). Transcription of pro-inflammatory immune mediators in the kidney (CxCL1/KC, CXCL2/MIP-2, CCL5/RANTES, TNFa and IFNg) were significantly more enriched in infected (TD and IP) than uninfected mice (OM and controls). This was also consistent with our previous observations for CJ (18). Others have done comparative analysis between susceptible and resistant strains of mice and have shown that levels of CxC chemokines are even more increased in mice that are resistant to infection (20). Transcription of the inflammatory and fibrosis markers in kidney (iNOS and ColA1) was equivalent to the uninfected controls, which was different than what we determined previously for IP and CJ infections (15), (18). Others have found that the loss of the iNOS gene had a negligible effect on the outcome of Leptospira infection, although they observed less susceptibility to nephritis in groups of mice lacking this gene (21). These results suggest that tracking expression of immunomediators in kidney and other tissues of strains of mice which are susceptible and resistant to infection helps shed light on Leptospira pathogenesis.

The B and T cell responses to *L. interrogans* were measured in blood and spleen of infected mice after transdermal (TD) and intraperitoneal (IP) exposure with the same dose of 10^8^ LIC (Fig. 4) and we found that the amount of IgM and IgG as well as the IgG subtypes enriched in the serum (IgG1) were similar to mice infected with lower doses of LIC via intraperitoneal route (15) and via the conjunctiva (18). Similar results were observed for the percentage of CD4+ and CD8+ T cell populations in spleen marked by a decrease of naïve cells which were activated in the presence of *Leptospira* and increased in the effector pool, whereas the number of memory T cells was not different from controls (15), (19). The same breakdown in populations of T cells is also observed in human patients (22), (23).

In summary, we found that transdermal (TD) infection leads to a delayed bacterial dissemination in blood and urine when compared to IP infection using the same dose, and that the overlapping presence of *L. interrogans* in both fluids was twice as long as the standard IP infection. Furthermore, dissemination of *L. interrogans* did not occur in mice infected with the same dose via the oral mucosa. Our findings underline the importance of precise determination of windows of pathogen dissemination in biological fluids and how the route of infection affects the outcome of disease progression. These studies are important to establish appropriate windows for application of direct or indirect diagnostic assays.

## ACKNOWLEDGMENTS

This work was supported by Public Health Service grants R44 AI096551 and R43 AI136551 (to M.G.S.) from the National Institutes of Health.

---

**Figure S1.**
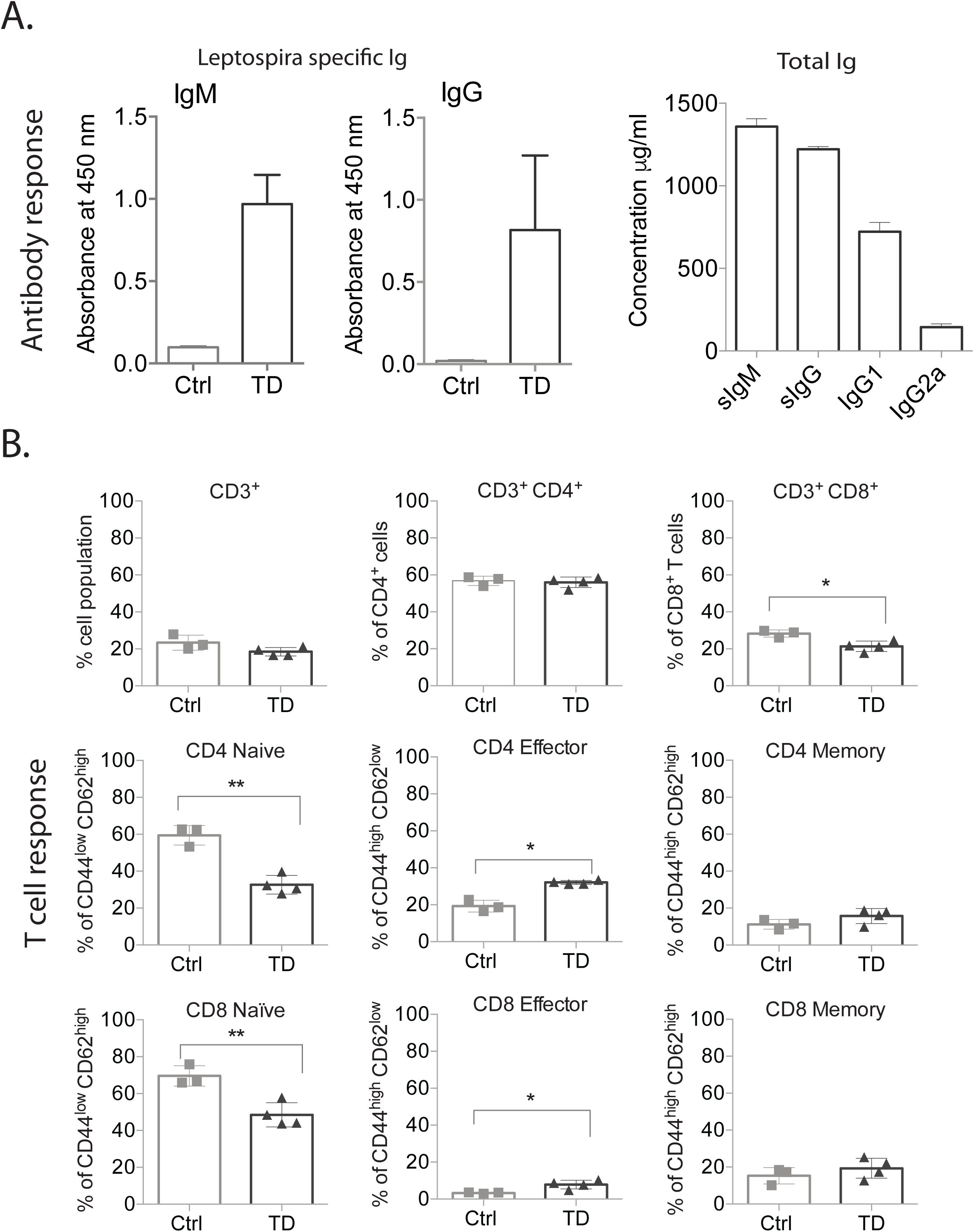
Immune response after *L. interrogans* infection. A. Leptospira-specific and total concentration of IgM and IgG in serum from mice infected transdermally with 10^8^ *L. interrogans*; IgG1 and IgG2a isotypes were also quantified by ELISA. B. Percentage of CD3+ T cell populations (CD4+, CD8+, naive, effector and memory) in spleen from infected mice and respective controls. Statistics by unpaired T test with Welch’ correction, *p<0.05 and ** p<0.005. N=3 or 4 mice per group. Data represents one of two independent experiments.

